# Changes in alpha activity reveal that social opinion modulates attention allocation during face processing

**DOI:** 10.1101/191916

**Authors:** Evelien Heyselaar, Ali Mazaheri, Peter Hagoort, Katrien Segaert

## Abstract

Participants’ performance differs when conducting a task in the presence of a secondary individual, moreover the opinion the participant has of this individual also plays a role. Using EEG, we investigated how previous interactions with, and evaluations of, an avatar in virtual reality subsequently influenced attentional allocation to the face of that avatar. We focused on changes in the alpha activity as an index of attentional allocation. We found that the onset of an avatar’s face whom the participant had developed a rapport with induced greater alpha suppression. This suggests greater attentional resources allocated to the interacted-with avatars. The evaluative ratings of the avatar induced a U-shaped change in alpha suppression, such that participants paid most attention when the avatar was rated as average. These results suggest that attentional allocation is thus an important element of how behaviour is altered in the presence of a secondary individual.

## 1. Introduction

A commonly observed phenomenon in psychological research is that an individual’s behaviour in a task is altered when conducted in the presence of another individual compared to when the task is done alone (Ringelmann, 1913). This effect is seen in a wide range of behaviours, for example participants eat more in the presence of someone else who is eating (Herman, 2015), participants’ performance in cognitive tasks decreases for complex tasks in the presence of a secondary individual (Bond & Titus, 1983), and these changes occur even when interacting with human-like computers instead of actual secondary persons (Mandell, Smith, Martini, Shaw, & Wiese, 2015). Although the effect of the presence of a secondary individual is not consistently positive or negative, there have been many behavioural studies establishing this phenomenon in different contexts. However, few have also investigated the mechanisms underlying the behaviour seen. Our study will be a step towards understanding how this process is implemented neurally.

In addition to the mere presence of a secondary individual, the opinion the participant has of this individual also influences the participants’ subsequent behaviour (Heyselaar, Hagoort, & Segaert, 2017; Lott & Lott, 1961; Weatherholtz, Campbell-Kibler, & Jaeger, 2014). One possible explanation for the phenomena discussed thus far involves the capture of attention by the secondary individual. Specifically, conducting a task in the presence of another person could cause one to divide their attention between the individual and the task, compared to when the task is done alone (for a review see Strauss, 2002). Additionally, if one finds the secondary individual more likeable, this could influence how much (or how little) attention is allocated to the secondary individual.

One way to investigate the neural processes related to attention is through electroencephalography (EEG), a non-invasive neuroimaging technique that measures the electrical potential generated by neurons. The EEG signal contains oscillatory activity in distinct frequency bands that have been found to map on to different facets of cognition (Siegel, Donner, & Engel, 2012). Alpha activity, an oscillation occurring at a frequency of 10 Hz, has been suggested to play a pivotal role in attention (Foxe, Simpson, & Ahlfors, 1998; Mazaheri & Jensen, 2010). According to this framework, the suppression of alpha activity relates to the degree of cortical activation whereas an increase in alpha activity relates to cortical inhibition. For this study we specifically focus on alpha activity as a representation of attentional allocation towards the secondary individual and modulations in the degree of alpha power/attention as a function of the opinion participants have of this secondary individual.

Here we have participants interact with and subsequently evaluate digital secondary individuals (hereafter “avatars”). We will measure the EEG activity during the viewing of the face of the avatars prior to their interaction and evaluation in Virtual Reality as well as after. As a control we will also present the face of avatars participants did not interact with. This design insures that the visual stimulus (in this case the face of the avatars) is constant, allowing us to investigate modulations of neural activity brought about by the interaction with, and evaluation of the avatars.

In sum, the first aim of our study is to determine whether viewing an avatar the participant has interacted with results in a different degree of attentional allocation (as indexed by changes in alpha activity) compared to viewing an avatar the participant has not interacted with. We will therefore provide neuroimaging evidence which may clarify why participant behaviour is different when conducting a task in the presence of a secondary individual. The second aim is to determine whether the amount of attention allocated (again measured as modulations in alpha activity) to the avatar varies as a function of the participants’ opinion of that avatar. Thus here we endeavour to provide a neurobiological explanation for behaviour seen in social psychology and social psycholinguistics.

## 2. Methods

### 2.1 Subjects

30 native Dutch speakers (2 male, M_Age_: 21.53 years, SD_Age_: 2.60) gave written informed consent prior to the experiment and were monetarily compensated for their participation. As the EEG cap had to fit underneath the virtual reality (VR) helmet, we were limited to testing participants with small head sizes (58cm diameter and below), restricting us to mostly female participants.

The previous study that used these avatars to observe an effect of opinion on behaviour (Heyselaar et al., 2017) showed an effect with 36 participants. However, as EEG effects are usually seen with a smaller sample size compared to behavioural studies, we decided to use 30 participants.

### 2.2 Procedure

The participants were informed that there were three phases to the experiment, but at the beginning of the experiment, they only received detailed information about Phase One. At the start of Phase Two they were informed of the goal of the study. The entire procedure is summarized in Figure 1.

**Figure 1.**
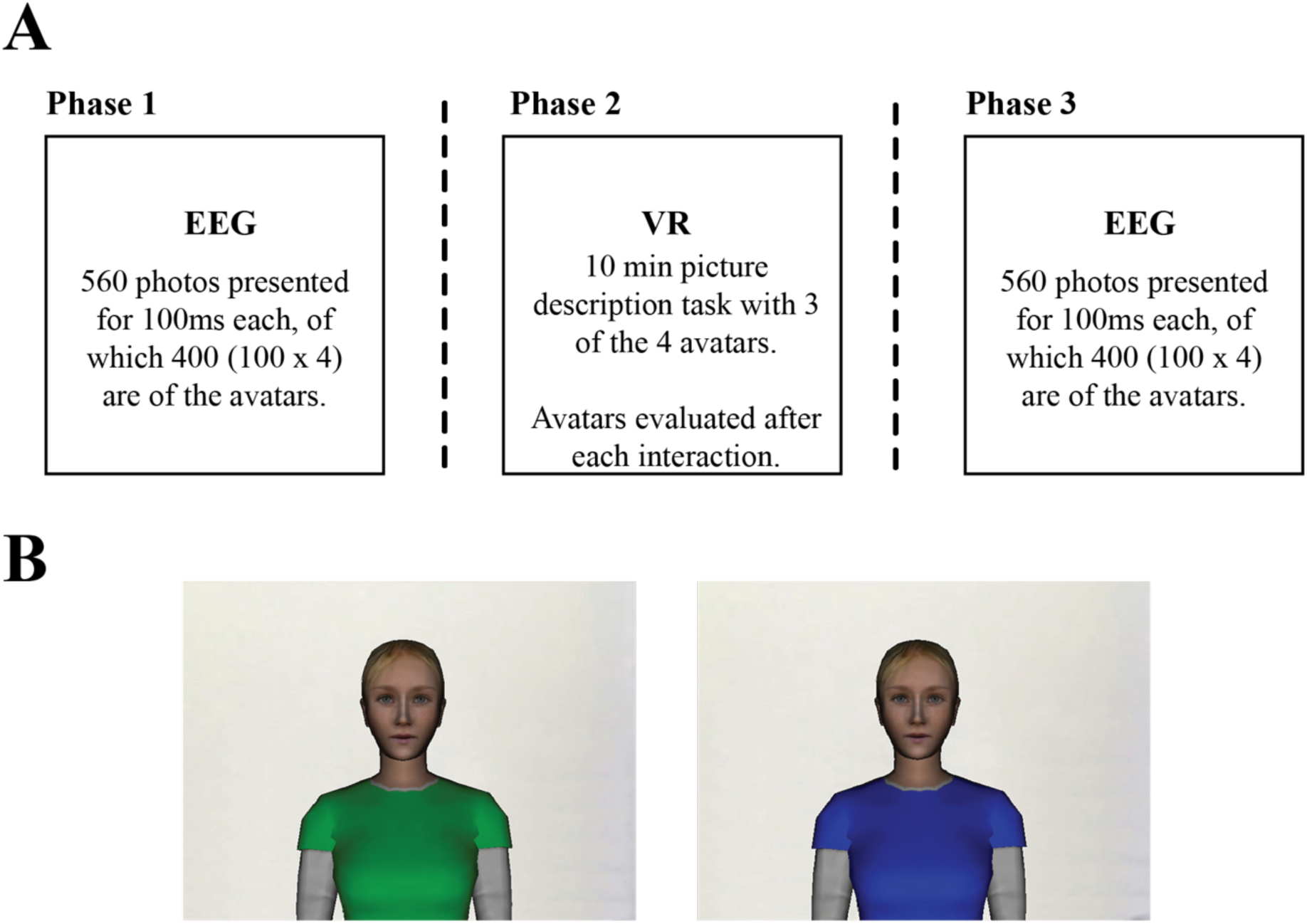
A. Procedure B. Examples of two of the four avatars. Avatars were presented with green, blue, red, and yellow shirts.

Participants initially viewed 560 static photos, of which 400 were of the 4 avatars. The faces of the four avatars were all exactly the same, and hence to be able to discriminate between photos of them, they were given different shirt colours. For Phase Two, participants interacted with three of the four avatars in Virtual Reality. Here the avatars were animated such that each avatar had different facial expressions in terms of their smile habit, eyebrow movements, and blink rate (see Table 1). Participants were therefore able to form opinions about the three avatars based on these facial characteristics, an effect seen robustly in previous studies using the same avatars and same facial expressions (Heyselaar et al., 2017). After each interaction, participants were asked to evaluate the avatars. For Phase Three, the participants were again shown static pictures. The participant used the colour of the shirt to discriminate between the avatars they had interacted with (and formed an opinion of) and the ones they had not interacted with. Details of each phase are given below:

#### 2.2.1 Phase One – Picture Evaluation Task

In Phase One participants viewed photos of avatars and filler pictures, with the aim to measure EEG responses of the participants to the avatar pictures before they had formed any opinion of these avatars. The participants were instructed to evaluate pictures as either “likeable” or “not likeable.” Each trial started with the presentation of a fixation cross for a 400 – 600ms jittered interval. This was followed by the presentation of a picture in the center of the screen. Pictures were presented for 100ms, followed by a jittered interval of 2000 – 3000ms before an evaluation screen was presented. Participants were asked to indicate, via a button press, whether they liked or disliked the picture they just saw. The location of the options (left or right for “likeable”) was randomized between participants.

Phase One consisted of 560 pictures. 400 of these pictures were of the avatars (100 repetitions of each avatar); the avatars are described below (*Materials - Avatars*). The remaining 160 were pictures selected from the pictures described below (*Materials - Picture Evaluation Task*). Picture order was randomized per participant.

#### 2.2.2 Phase Two – Picture Description Task

Participants interacted with three avatars for 10 minutes each in the virtual environment (these avatars are hereafter referred to as “Interacted-with avatars”). The EEG cap remained on in Phase Two, but no activity was recorded. The fourth avatar was not interacted with as a control (hereafter “Non-interacted-with avatars”). The shirt colour of all the avatars (both Interacted-with and Non-interacted-with) was pseudo-randomized across participants.

Participants at this point were informed of the goal of the study, to ensure they paid as much attention to the relevant characteristics of the avatar as possible.

The participant and the avatar would alternate in describing picture cards to each other. If the listener saw the card described by their partner as one of the cards in their spread they would select it, causing it to be automatically replaced by a novel card. The listener would then become the speaker and pick a card to describe. This continued until 50 cards were described, after which the headset was removed and participants were asked to fill out a pen-and-paper questionnaire about the avatar. We favoured a pen-and-paper questionnaire instead of having the avatar ask the questions directly as previous research has shown that if the participant evaluates the avatar in the presence of said avatar, they rate them more favourably (Nass, Moon, & Carney, 1999).

The questionnaire consisted of three 6-point Likert-scale questions asking to rate the avatar on perceived humanness, perceived strangeness, and quality of their facial expressions in relation to the other two avatars. After each avatar, the participants were allowed to change their ratings for previously viewed avatars. The order of the avatars was pseudo randomized across participants.

#### 2.2.3 Phase Three – Picture Evaluation Task

In Phase Three, we again recorded EEG activity while participants viewed photos of the four avatars mixed with filler pictures. We kept Phase Three the same as Phase One, using the same picture order, therefore any difference in the modulation of alpha activity induced by viewing pictures of the avatars between Phase One and Phase Two would likely be due to the social interaction that occurred in VR in Phase Two. By comparing the Interacted-with Avatar to the Non-interacted-with Avatar we can control for any repetition effects as both are viewed in both Phases. Before EEG recordings in Phase Three began, impedance for each electrode was checked and adjusted as necessary.

### 2.3 Materials

#### 2.3.1 Picture Evaluation Task (Phase One and Phase Three)

165 pictures were taken from the Geneva Affective PicturE Database (GAPED; Dan-glauser & Scherer, 2011). An equal number (55) belonged to the category positive, negative or neutrally-rated pictures, in terms of emotional valence. We attempted to ensure that the arousal rating was comparable between picture categories as much as possible. The average ratings for arousal were thus 30.47 (SD: 9.491), 29.52 (SD: 5.92), and 46.46 (SD: 7.01) for positive, neutral, and negatively rated pictures respectively.

We also included four avatar photos: the same picture of the avatar with a yellow, green, red or blue shirt.

#### 2.3.2 Avatars and Virtual Environment in Phase Two

All avatars had the same exterior adapted from a stock avatar produced by WorldViz (“casual15_f_highpoly”; see Figure 1B). All the avatars’ speech was pre-recorded by the same human female and played during appropriate sections of the experiment. The avatars’ appearance suggested that she was a Caucasian female in her mid-twenties, which matched the age and ethnicity of the Dutch speaker who recorded her speech.

The three facial expressions used have been tested elsewhere and have been convincingly demonstrated to induce a wide spread of ratings with regard to perceived humanness, perceived strangeness, and quality of facial expression (Heyselaar et al., 2017). These three facial expressions involved combinations of subtle changes in blink rate, smiling, and eyebrow habits (Table 1). Facial expression choices were based on work done by Looser & Wheatley (2010) who have shown that perception of humanness is dependent on upper face movement.

Blinks happened once every 1 - 5 seconds. For versions with normal smiling and normal eyebrow habits we explicitly programmed when the avatar would smile and/or raise her eyebrows, such that it would coincide with the content of her speech. For example, the avatar would raise her eyebrows when asking a question and smile when she was enthusiastic. When not speaking, she would smile once every 5 - 10 seconds and raise her eyebrows once every 1 - 5 seconds such that she would still differ from the no smile/no eyebrow version. All of these changes were extremely subtle to ensure that they can still be related to ecologically valid behavioural characteristics that one would encounter in the everyday world.

**Table 1.** Avatar Facial Expressions.

| Avatar | Blink Duration<sup>1</sup> | Smiling Habit | Eyebrow Habit |
| --- | --- | --- | --- |
| 1 | No blink | No smile | No movement |
| 2 | 0.1 sec (Normal) | Dialogue-matched | Once every 3 - 5 sec |
| 3 | 0.1 sec (Normal) | Dialogue-matched | Dialogue-matched |

<sup>1</sup>Measured from the beginning of the closing movement to when the eye is fully open again

The virtual environment (VE) was a stock environment produced by WorldViz (“room.wrl”) adapted to include a table with a wooden divider. We chose to have the cards displayed at the top of the divider so that the participants could see the cards while facing forward. This was done due to the weight of the head-mounted display (HMD), which would cause an uncomfortable strain on the back of the participants’ heads when they face down. Having the participants face forward throughout the entire experiment distributes this weight more comfortably.

The experiment was programmed and run using WorldViz’s Vizard software. Participants wore an NVIS nVisor SX60 HMD, which presented the VE at 1280 × 1024 resolution with a 60-degree monocular field of view. Mounted on the HMD was a set of 8 reflective markers linked to a passive infrared DTrack 2 motion tracking system from ART Tracking, the data from which was used to update the participant’s viewpoint as she moved her head. It is known that this type of headset can cause dizziness and nausea due to the exclusion of the participant’s nose in the field of view (Whittinghill, Ziegler, Moore, & Case, 2015). However, as each interaction was quite short (~5 minutes), none of our participants reported feeling any nausea.

Additionally, a single reflective marker was taped onto the index finger of the participant’s dominant hand. This marker was rendered as a white ball in the VE, such that participants knew the position of their finger at all times. Sounds in the VE, including the voice of the avatars, were rendered with a 24-channel WorldViz Ambisonic Auralizer System.

#### 2.3.3 Picture Description Task in Phase Two

The pictures used in this task have been described elsewhere (Segaert, Menenti, Weber, & Hagoort, 2011). Our stimulus pictures depicted 40 transitive events such as *kissing, helping* or *strangling* with the agent and patient of this action. Each event was depicted by a greyscale photo containing either one pair of adults or one pair of children. These pictures were used to elicit transitive sentences; for each picture speakers can either produce an active transitive sentence (e.g. *the woman kisses the man*) or a passive transitive sentence (e.g. *the man is kissed by the woman*).

We also included pictures depicting intransitive events such as *running, singing,* or *bowing* using one actor. The actor could be any of the actors used in the transitive stimulus pictures. Each card consisted of one stimulus picture with the relevant verb printed underneath.

### 2.4 Data Analysis Approach

#### 2.4.1 Pre-processing

EEG was recorded from 64 cap-mounted Ag/AgCl electrodes (ActiCAP, Brainproducts). Horizontal eye movements were monitored by two electrodes placed at the outer left and right canthi. Vertical eye movements were monitored using an electrode placed below the left eye. In addition, electrodes were placed on the left and right mastoid bones. During EEG recording, all electrodes were referenced to the left mastoid. All impedances were kept below 10kΩ. Signals were recorded with a BrainAmp amplifier system, using a 150 Hz low-pass filter, a time constant of 10s (0.016 Hz), and a 500 Hz sampling rate. Signals were later re-referenced off-line to linked mastoids.

The pre-processing of the data was done using functions from EEGLAB (Delorme & Makeig, 2004) and the Fieldtrip software package (Oostenveld, Fries, Maris, & Schoffelen, 2011). Fieldtrip EEG epochs were locked to the onset of the picture and manually inspected for non-physiological artefacts. Ocular artifacts were removed using independent component analysis (infomax algorithm) incorporated as the default “runica” function in EEGLAB.

#### 2.4.2 Time-Frequency Representations (TFR) of Power

Time-frequency representations (TFR) of power were calculated for each trial using sliding Hanning tapers having an adaptive time window of three cycles for each frequency of interest (Δ*T* = 3/*f*), utilizing the same approach used in previous studies e.g., Mazaheri et al. (2014) and van Diepen, Cohen, Denys, & Mazaheri (2015).

The classification of frequency bands in delta (2 – 4 Hz), theta (5 – 8 Hz), alpha (8 – 14 Hz), high alpha (11 – 14 Hz), and beta (15 – 20 Hz) for further analysis were based on prior literature (Hamel-Thibault, Thénault, Whittingstall, & Bernier, 2015; Kliegl, Pastötter, & Bäuml, 2015; Klimesch, 1997; Lange, Oostenveld, & Fries, 2013; Shahin & Pitt, 2012).

Changes in the power of oscillatory activity induced by the onset of avatar faces were expressed in terms of change scores from baseline (Δ*P*) using the following formula: Δ*P* = (*Pt* − *Pr*)/*Pr*, where *Pr* was the mean power during the baseline period 150ms to 650ms before the onset of the picture and *P*t was the power at each specific time point.

### 2.5 Statistical Analysis

#### 2.5.1 Correction for multiple comparisons

We corrected for multiple-comparisons (multiple electrodes) by means of a non-parametric cluster level (over-electrodes) randomization routine (Maris & Oostenveld, 2007). In this procedure, for each contrast (e.g., Phase 3 versus Phase 1) first a two-tailed dependent *t*-test was computed for each individual electrode-time pairs. Next, electrodes which passed the significance threshold at a 5% significance level were clustered by direction of effect and spatial proximity. A Monte Carlo estimated probability value was estimated for this cluster by randomly swapping the conditions within subjects and calculating the maximum cluster-level test statistic a 1,000 times. We have employed a similar procedure coded in the Fieldtrip toolbox in a number of previous studies (van Diepen et al., 2015; van Diepen, Miller, Mazaheri, & Geng, 2016; van Diepen & Mazaheri, 2017).

#### 2.5.2 Mixed models

The values extracted from the electrodes of interest (see *Results*) were analysed using a linear mixed effects model, using the lmer function of the lme4 package (version 1.1.9; Bates, Maechler, & Bolker, 2012) in R (R Core Development Team, 2011). The dependent measure was the values extracted from the regions of interest. The repeated-measures nature of the data was modelled by including a per-participant and per-shirt-colour random adjustment to the fixed intercept (“random intercept”). We began with a full model (two-way interactions between each of the ratings) and then performed a step-wise “best-path” reduction procedure, removing interactions before main effects, to locate the simplest model that did not differ significantly from the full model in terms of variance explained. All ratings were centred before entry into the model. *P* values were extracted using the Anova function from the car package (version 2.1.0; Fox & Weisberg, 2011) using Wald Chi-Square tests (Type III).

## 3. Results

All data were normalized as percent change from a baseline interval (150 – 650ms before picture onset) within participants to reduce the contribution of participants with large variance in the power estimates. To identify time windows of interest we compared the time-frequency data between Phase 3 and Phase 1 (Figure 2A). These data represent changes in spectral power due repetition effects, as the task was exactly the same between Phase 1 and Phase 3. A non-parametric cluster-based permutation analysis was used to determine time points of interest for delta, alpha, and beta frequency bands. Time points of interest identified are listed in Table 2, and illustrated in Figure 2B.

**Table 2.** Time points of significant difference in oscillatory power between Phase 3 and Phase 1. *P* values were obtained using Monte Carlo simulations (1,000 iterations), corrected for multiple comparisons.

| Frequency Range (Hz) | Time Period (ms) | *p* value |
| --- | --- | --- |
| 2 – 4 (Delta) | 900 – 1200 | .036 |
| 5 – 8 (Theta) | 100 – 350 | two clusters; $p < .020$ |
|  | 700 – 1500 | .003 |
| 8 – 14 (Alpha) | 100 – 500 | two clusters; $p < .047$ |
| | 750 - 1000 | two clusters; $p < .006$ |
| 15 – 20 (Beta) | 700 – 1150 | .002 |

**Figure 2.**
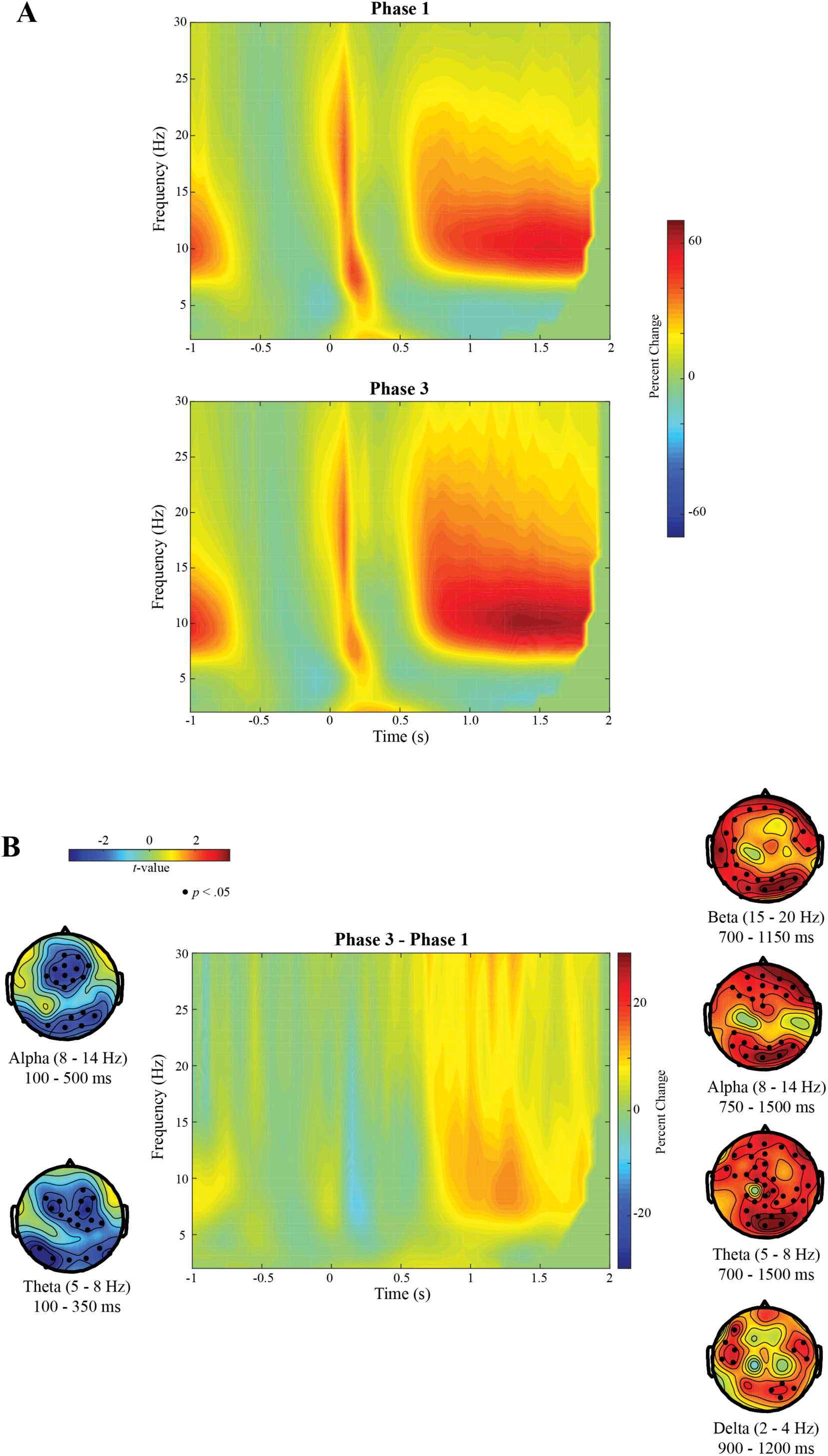
TFR of all conditions for all electrodes for A. Phase 1 and Phase 3 separately and B. for Phase 3 minus Phase 1. We used a non-parametric cluster based permutation test to identify time points of interest that were significantly different between Phase 3 and Phase 1. These time intervals of interest (TOI) are used in the subsequent analysis. The top panel of section B shows the topography of the significant power changes in the different frequency bands. The electrodes which showed a significant different between Phase 3 and Phase 1 are marked with dots (*p* < .05, Monte Carlo estimated). TFRs are expressed as a percentage change from baseline (−650ms to −150ms before face onset).

To control for low-level sensory-induced oscillatory changes in the EEG induced during the onset of pictures, we subtracted the EEG activity in Phase One from the EEG activity in Phase Three. This ensures that we are subtracting any EEG activity related to visual onset of the pictures (and performing the ‘like/dislike’ task) since this occurred in Phase One as well as in Phase Three.

### 3.1 Interacted-with avatars induced greater alpha suppression than Non-interacted-with avatars

We set out to investigate whether oscillatory activity is modulated by the experience of interacting with and evaluating the traits (both occurring in Phase 2) of 3 of the 4 avatars. We limited our analysis to the time and frequency windows identified above.

The comparison of alpha power between Interacted-with and Non-interacted-with avatars revealed that for the Interacted-with avatars high alpha (11 – 14 Hz) was significantly more suppressed 400 – 450ms post-face onset for a cluster of parietal electrodes (Monte Carlo *p* = .017, corrected for multiple comparisons). Later, alpha activity (8 – 14 Hz) between 750 – 1000ms post-face onset (Monte Carlo *p* = .050, corrected for multiple comparisons) was also significantly more suppressed for Interacted-with. There was also a significantly greater suppression of beta power 700 – 1000ms post-face onset (Monte Carlo *p* = .015, correct for multiple comparisons) for Interacted-with compared to Non-interacted-with avatars for a cluster of midline central electrodes. There were no significant effects found in any of the other time intervals of interest. The significant clusters are illustrated in Figure 3.

**Figure 3.**
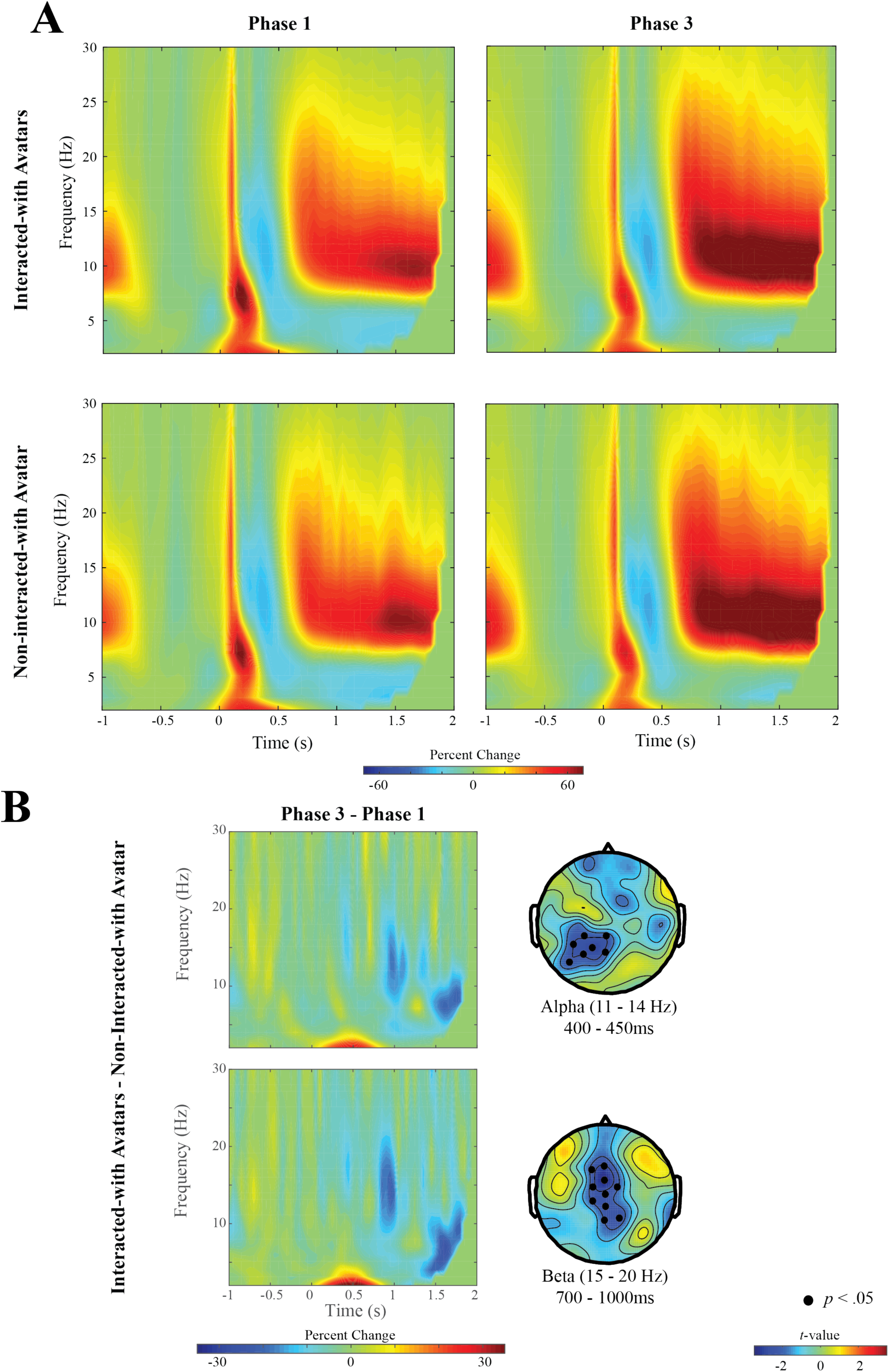
Power difference between the Interacted-with and Non-interacted-with avatars. **A.** The TFR of all electrodes for the Interacted-with and Non-interacted-with avatars for Phase 1 and Phase 3. **B.** The TFR for the cluster of electrodes showing a significant difference between Interacted-with avatars and Non-interacted-with avatars, after subtracting Phase One from Phase Three. This comparison reflects processing related only to having interacted with the avatar and evaluating their traits (both occurring in Phase Two). The topography contains the two significant clusters (marked with dots): 11 – 14Hz (Alpha), 400 – 450ms post-face onset; and 15 – 20Hz (Beta), 700 – 1000ms post-face onset. TFRs are expressed as a percentage change from baseline (−650ms to −150ms before face onset).

### 3.2 Amount of alpha suppression for the Interacted-with avatars was modulated by social opinion ratings

The increase in alpha suppression observed for Interacted-with avatars suggests greater attentional resources were allocated to them than to the Non-interacted-with avatars. The second aim of this study is to determine whether the alpha modulation (i.e. attention allocation) varied as a function of how the avatars were rated on a certain trait.

The purpose of having the participants interact with three different avatars was to ensure as wide a range of ratings per trait as possible. Table 3 shows the spread of ratings for each of the three traits (perceived humanness, perceived strangeness, and quality of facial expression) where a higher rating represents a higher score in said trait (more human, more strange, higher quality of facial expression). Participants were asked after interacting with each avatar to rate said avatar on each of these traits *in relation to the other two avatars.* For this reason, participants were allowed to change their answers after having interacted with all three.

**Table 3.** Number of Ratings in Each Likert-Scale Trait. Three traits were tested (perceived humanness, perceived strangeness and quality of facial expressions). Higher ratings represent a higher score in said trait (more human, more strange, higher quality of facial expression).

| Trait | Ratings |  |  |  |  |  |
| --- | --- | --- | --- | --- | --- | --- |
|  | 1 | 2 | 3 | 4 | 5 | 6 |
| Perceived Humanness | 1 | 18 | 22 | 37 | 10 | 2 |
| Perceived Strangeness | 7 | 34 | 25 | 21 | 3 | 0 |
| Quality of Facial Expression | 9 | 25 | 21 | 24 | 9 | 2 |

As predicted, we have a wide spread of ratings across most of the traits, with only the outer most ratings having 3 data points or less. Before using these data in any analysis, they were first trimmed to only include ratings with 7 or more data points. In other words, we removed rating 1 (N = 1) and 6 (N = 2) for perceived humanness, rating 5 (N = 3) and 6 (N = 0) for perceived strangeness, and rating 6 (N = 2) for quality of facial expression.

To determine if these ratings can modulate the activity shown in Figure 3, we used the channels that showed significant activity as regions of interest. We extracted the average values for each participant for each avatar for the alpha, beta, and theta band frequency and time windows as specified above. The values for Phase 1 were then subtracted from the values for Phase 3 to control for any changes in activity related to the processing of photos. In the resulting dataset, the values for the Non-interacted-with avatars were then subtracted from the values for the Interacted-with avatars to control for the fact that the photos were viewed for a second time. The resulting values therefore capture any changes in activity due to how participants process the Interacted-with avatars. These resulting values were then entered into a mixed model along with the trimmed behavioural ratings for each of the three traits in Table 2.

The model for the picture-induced alpha suppression (400 – 450ms; 11 – 14 Hz) showed a tendency for the Perceived Humanness rating to predict alpha activity (χ^2^(1) = 3.55, *p* = .059). Perceived Strangeness was in the best model, but showed no significant contribution (χ^2^(1) = 0.20, *p* > .250). Figure 4 shows the influence of Perceived Strangeness and Perceived Humanness on changes in alpha activity between Interacted-with and Non-interacted-with avatars.

**Figure 4.**
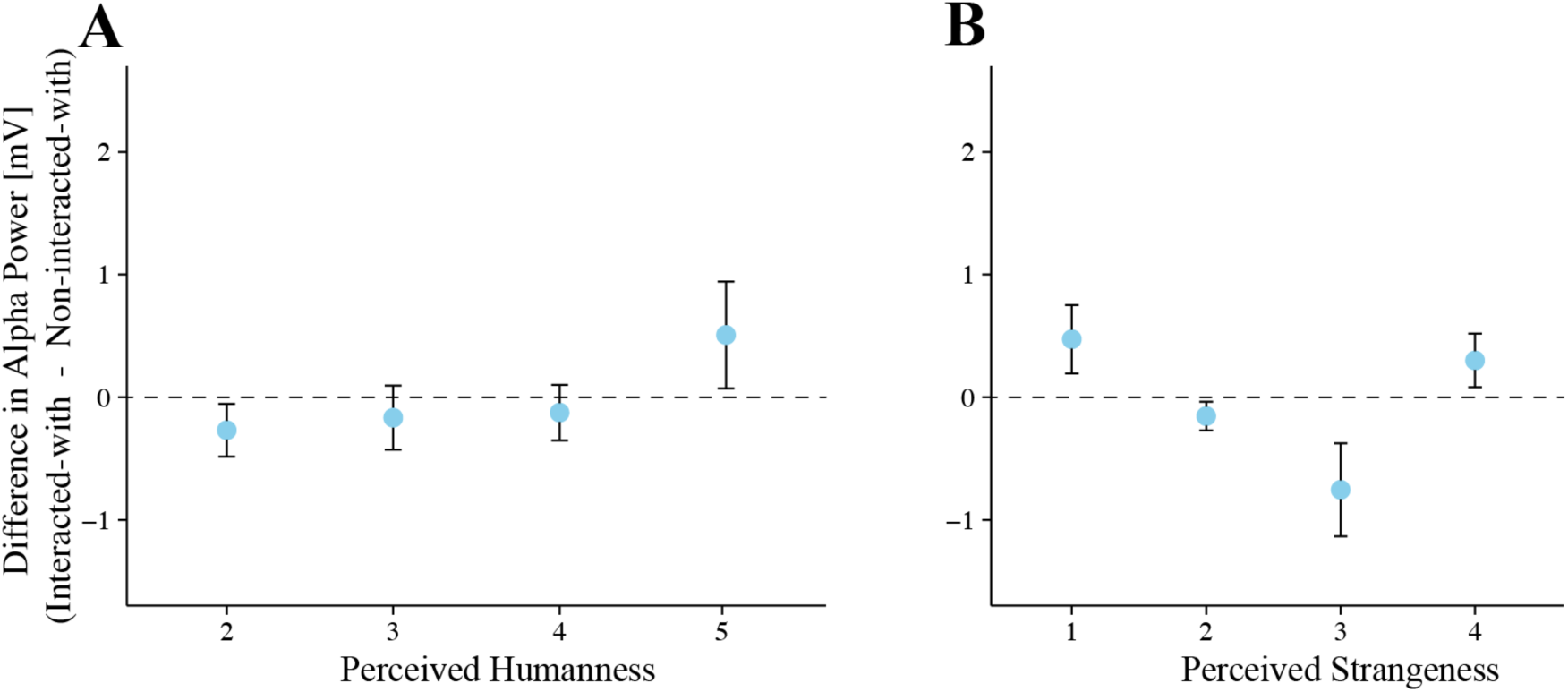
Difference in Alpha Activity between Perceived Humanness (A) and Perceived Strangeness (B) Ratings. We only included ratings with more than 7 datapoints in the model. These ratings are illustrated here. Only ratings for perceived strangeness (modelled as a quadratic term) significantly predicted the difference in alpha power to the presentation of the faces of the Interacted-with avatar compared to the Non-interacted-with avatar. This suggests that the medially-strange rated avatar (rating of 3 out of 6) was allocated the most attention.

Due to the quadratic shape of the effect of Perceived Strangeness on alpha activity, we re-ran the mixed effects model with Perceived Strangeness modelled as a quadratic term. The results of this model are summarized in Table 4. The quadratic term significantly predicted the alpha suppression (*p* = .010). This result suggests that perception of this trait modulates attention allocation such that avatars which are perceived as being very strange or not very strange draw the least amount of attention, whereas those that are perceived as being medially-strange draw the most amount of attention.

**Table 4.** Summary of best linear mixed model for changes in alpha activity between Interacted-with and Non-interacted-with avatars.

|  | coefficient | SE | t value | p value |  |
| --- | --- | --- | --- | --- | --- |
| Intercept | -0.41 | 0.17 | -2.49 | .010 | * |
| Perceived Strangeness (quad.) | 0.10 | 0.04 | 2.56 | .010 | * |
| Perceived Humanness (linear) | -0.21 | 0.12 | -1.78 | .075 | . |

N = 84

Even though our main interest was in modulations of alpha suppression, which represent modulations in attention allocation, as we found significant changes in beta power between the Interacted-with and Non-interacted-with avatars, we next also modelled these changes in the same way as described above.

The model for picture-induced beta suppression (700 – 1000ms; 15 – 20 Hz) showed a tendency for the Quality of Facial Expression ratings to predict beta activity. Beta oscillations are commonly correlated with imagined motor preparation (McFarland et al., 2000; Pfurtscheller et al., 1997) and hence these correlations may be indicators of what factors the participants use to rate the evaluated avatars in Phase 3. Beta values of the model showed that increased ratings of Quality of Facial Expression decreased (β = −0.10) beta suppression in the Interacted-with compared to the Non-interacted-with avatar.

## 4. Discussion

It is well established in psychological science that the presence of a secondary individual influences a participants’ behaviour on a task and, moreover, the opinion the participant has of this secondary individual also affects behaviour. We hypothesized both phenomena can be explained by the capture of attention. In the current study we investigated this by examining changes in alpha suppression after participants had interacted with and evaluated digital individuals (“avatars”). The first aim of our study was therefore to determine whether viewing an avatar the participant has interacted with resulted in a different degree of attentional allocation and we found that this was indeed the case. We also examined if the changes in alpha activity observed differed as a function of the ratings participants had given the secondary individual before we recorded their EEG activity and indeed found a relationship between evaluation ratings of the secondary individual and modulations in alpha suppression. We therefore conclude that attentional capture, as measured by changes in alpha activity, is an important step towards understanding how secondary individuals influence participants’ behaviour.

We will go into the results of each of our two aims in detail. Our first aim was to determine whether attentional resources were differentially allocated when participants viewed faces of individuals they had just interacted with. To test this, participants viewed pictures of avatars before and after interacting with and evaluating them (Interacted-with avatars). While viewing the avatar’s faces, we recorded participants’ EEG activity. The Non-interacted-with avatars appeared in both parts of the EEG experiment but were not interacted with. We initially identified four frequency bands of interest: delta, theta, alpha, and beta. Only the alpha and beta bands showed a significant difference when participants viewed faces of Interacted-with compared to Non-interacted-with avatars.

We found that the Interacted-with avatars induced a greater amount of alpha suppression 400ms after picture onset compared to the Non-interacted-with avatar. Alpha suppression occurred over the occipital parietal cortices with a high peak frequency (11 – 14 Hz), which has been previously associated with semantic processing demands (Klimesch, 1997). This suggests that pictures of the Interacted-with avatars have been allocated more resources for processing compared to the Non-interacted-with avatars. The left posterior parietal location of this cluster is suggestive of an effect indicating enhanced perceptual analysis of the input (De Cesarei & Codispoti, 2011).

We hypothesize that the beta suppression seen 700 – 1000ms post stimulus onset is indicative of what factors influences the participant’s motor response in the ‘Like/Dislike’ task that followed 2 seconds post stimulus onset (Pfurtscheller et al., 1997; McFarland et al., 2000). The central location of the significant clusters as opposed to the ipsi-/contra-lateral locations usually seen in the literature is most likely due to us randomizing the location of the Like and Dislike button between participants.

Our second aim was to determine whether the degree of picture-induced alpha modulation for Interacted-with avatars differed as a function of the opinion the participant had of the secondary individual. All the avatars were exactly the same, except for the facial expressions, which we have shown in two previous studies to be enough to vary the opinion of these avatars (Heyselaar et al., 2017; Heyselaar et al., 2015). However, as they were identical in appearance, we cannot rule out the possibility that they were perceived not as individuals but as the same avatar with varying moods. This does not change the interpretation of our results, as it is still the opinion one has of the secondary individual that effects attentional allocation.

We found a modulation of alpha suppression as a function of the perceived strangeness rating in the form of a U-shaped curve for the early alpha effect (400 – 450ms). The most- and least-strangely rated avatars induced less alpha suppression for the Interacted-with compared to the Non-interacted-with avatars, whereas the medially-rated avatars elicited less alpha suppression for the Non-interacted-with compared to the Interacted-with avatars. This suggests that the perception of the most- and least-strangely rated avatars requires fewer resources compared to perception of the medially-rated avatars.

The results of the current study are consistent with previous literature looking at the effect of face likeability and distinctiveness on memory performance. Lin and colleagues (2011) had participants learn adjectives associated with faces and observed a U-shaped, quadratic curve such that participants had worse recall performance for the adjectives matched with faces rated as averagely-distinctive and best performance for adjectives presented with likeable and unlikeable faces. Hence, the effect on attention that we observe in our study could be due to more resources being required to retrieve information about the medially-rated avatar, as they were not remembered as well as the most- and least-strangely rated avatars.

The avatars we used in this study have been used before to elicit a difference in syntactic priming as a function of the strangeness rating (Heyselaar et al., 2017). Syntactic priming refers to the phenomenon where participants increasingly use their partners’ grammar in their own utterances (Bock, 1986), and recent studies have shown that the magnitude with which participants adapt their language behaviour varies as a function of how the participant rated their partner. In our previous study, we observed an inverted U-shaped curve such that the maximal syntactic priming effect occurred for the middle ratings of perceived strangeness for the same avatars we used in the current study. To relate this to participant performance in the presence of a secondary individual, when avatar interaction happens in concurrence with a task, such as syntactic priming, the participant has to divide attention between the faces and the task. Therefore, when little attention is given to the avatar itself (as with the medially-rated avatars), then more attention is available to complete the task, which would result in a better performance (i.e., more syntactic priming) at this task compared to doing the task in the presence of a secondary individual the participant finds extremely likeable or unlikeable. This is in line with the distraction-conflict hypothesis (Baron, 1986): The presence of others is a distraction, which leads to attentional conflict in terms of cognitive overload and selective focusing of attention.

To conclude, the first aim of our study was to determine whether attentional allocation varies when viewing faces of an individual the participant has interacted with and evaluated compared to not. We are confident that the capture of attention plays a role in modulating the behavioural effects seen in the social psychology literature. Our second aim was to determine whether this allocation of attention differs as a function of rating and we show that it does. This clarifies why participant behaviour is different depending on whether they positively or negatively view the secondary individual, an effect seen not only in social psychology but also in psycholinguistics.

